# Nucleoporins facilitate ORC loading onto chromatin

**DOI:** 10.1101/2022.04.05.487200

**Authors:** Logan Richards, Christopher L. Lord, Mary Lauren Benton, John A. Capra, Jared T. Nordman

## Abstract

The origin recognition complex (ORC) binds throughout the genome to initiate DNA replication. In metazoans, it is still unclear how ORC is targeted to specific loci to facilitate helicase loading and replication initiation. Here, we performed immunoprecipitations coupled with mass spectrometry for ORC2 in Drosophila embryos. Surprisingly, we found that ORC2 associates with multiple subunits of the Nup107-160 subcomplex of the nuclear pore. Bioinformatic analysis revealed that, relative to all modENCODE factors, nucleoporins are among the most enriched factors at ORC2 binding sites. Critically, depletion of the nucleoporin Elys, a member of the Nup107-160 complex, results in decreased ORC2 loading onto chromatin. Depleting Elys also sensitized cells to replication fork stalling, which could reflect a defect in establishing dormant replication origins. Our work reveals a new connection between ORC, replication initiation and nucleoporins, highlighting a previously unrecognized function of nucleoporins in metazoan replication initiation.

## INTRODUCTION

The origin recognition complex (ORC) binds to thousands of sites throughout the genome to initiate DNA replication (Leonard and Méchali, 2013). Chromatin-bound ORC, together with additional factors, performs the essential function of loading MCM2-7 helicases in an inactive state across the genome in late M and G1 phases of the cell cycle (Fragkos et al., 2015). The distribution of ORC-binding sites is critical to define replication start sites and to maintain genome stability, as large genomic regions devoid of replication start sites are prone to breakage upon replication stress (Cha and Kleckner, 2002; Letessier et al., 2011; Miotto et al., 2016; Newman et al., 2013). Additionally, the number and distribution of ORC-binding sites and replication start sites can change during development to accommodate cell-type-specific DNA replication programs (Eaton et al., 2011; Hua et al., 2018; Sher et al., 2012). Therefore, understanding how ORC is targeted to chromatin is an important parameter in maintaining genome stability during development.

The factors that determine where ORC binds differ across species; however, both DNA sequence and chromatin environment can be important contributors. In *S. cerevisiae*, ORC binding is largely sequence dependent and highly influenced by nucleosome positioning (Eaton et al., 2010; Wyrick et al., 2001; Xu et al., 2006). While there are a small number of defined initiator sequences metazoans (Altman and Fanning, 2001; Austin et al., 1999; Lu et al., 2001), ORC binding is largely sequence independent and highly influenced by both chromatin state and DNA topology (Eaton et al., 2010, 2011; MacAlpine et al., 2010; Miotto et al., 2016; Remus et al., 2004; Vashee et al., 2003). ORC tends to localize to the transcription start sites of active genes (Eaton et al., 2011; MacAlpine et al., 2010). Hallmarks of ORC binding include open and accessible regions of chromatin, histone modifications associated with active chromatin such as H3K27ac and H3K4me3 and in Drosophila, sites of cohesion loading (Eaton et al., 2011; MacAlpine et al., 2010; Miotto et al., 2016). The relatively non-specific ORC binding in metazoans likely facilitates developmental-specific replication programs. Furthermore, in Drosophila, specific proteins such as E2f, Rbf and a Myb-containing protein complex can help recruit ORC to a specific initiation site (Beall et al., 2002; Bosco et al., 2001; Royzman et al., 1999). In humans, ORC-associated protein (ORCA) localizes to heterochromatin and facilitates ORC loading onto chromatin (Shen et al., 2010). The number of ORC-binding sites greatly exceeds the number of replication start sites used in a given cell cycle (Cayrou et al., 2011). These excess ORC-binding sites license dormant replication origins, which have a critical role in promoting genome stability by ensuring additional replication start sites are available upon replication stress (Doksani et al., 2009; Ge et al., 2007; Ibarra et al., 2008).

Nucleoporins, or Nups, are typically associated nuclear pore complexes (NPCs) and facilitate the import and export of proteins and macromolecules across the nuclear membrane (for review, see Wente and Rout, 2010). In addition to their canonical function at NPCs, a subset of Nups bind to chromatin and regulate genome structure and function. For example, the nucleoporin Elys binds to chromatin in late mitosis and is required to assemble nuclear pore complexes onto chromatin prior to their insertion into the nuclear membrane (Franz et al., 2007; Galy et al., 2006; Gillespie et al., 2007; Rasala et al., 2006; Shevelyov, 2020). More recent work; however, has demonstrated that several Nups regulate both transcription and chromatin condensation (Capelson et al., 2010; Kalverda et al., 2010; Kuhn et al., 2019; Panda et al., 2014; Pascual-Garcia et al., 2014, 2017; Raices and D’Angelo, 2017; Vaquerizas et al., 2010). In Drosophila, Nup98 binds to distinct regions of the genome, co-localizes with RNA polymerase II and regulates mRNA levels (Panda et al., 2014; Pascual-Garcia et al., 2014). Nup98 can also facilitate promoter-enhancer contacts at least at a subset of genes in Drosophila (Pascual-Garcia et al., 2017). Furthermore, the genomic localization of Nup98 and Elys correlate with actively transcribed genes (Pascual-Garcia et al., 2017). Tethering the nucleoporins Nup62 or Sec13 is sufficient to decondense chromatin within specific regions of chromatin (Kuhn et al., 2019). Interestingly, this chromatin decondensation correlates with the recruitment of Elys and the PBAP/Brm chromatin remodelling complex (Kuhn et al., 2019). Many Nups are not permanently anchored to the NPC, but, rather, dynamically associate with the NPC throughout the cell cycle (Rabut et al., 2004) and the interaction between Nups and chromatin can occur in the nucleoplasm (Ibarra and Hetzer, 2015). Therefore, it is likely that many Nups have chromatin-related functions independent of the NPC.

In this study, we show that ORC associates with members of the Nup107-160 subcomplex of the nuclear pore. We then show that Nups co-localize with ORC2-binding sites across the genome and that Nups are some of the most enriched chromatin-related factors at ORC sites. We find that depletion of Elys, but not other Nups, reduces the amount of chromatin-bound ORC2 throughout the genome. Importantly, Elys likely promotes ORC2 association independently of its role in promoting chromatin decompaction as we observe no difference in chromatin accessibility at ORC2 binding sites upon Elys depletion. Finally, we show that depletion of Elys and Nup98-96 sensitizes cells to replication fork inhibition. We propose that Elys is necessary to load the optimal level of ORC on chromatin. Reduction in ORC levels upon Elys depletion could underlie the sensitivity to replication fork stalling by jeopardizing the establishment of dormant origins. Thus, our work provides new insight into how metazoan ORC is recruited to chromatin and defines a replication-associated function of Nups in Drosophila.

## RESULTS

### ORC associates with nucleoporins

ORC binding throughout the genome is essential for initiation of DNA replication, genome stability and replication timing (Eaton et al., 2011; Leonard and Méchali, 2013; Miotto et al., 2016). While a number of chromatin-associated factors are important for ORC genomic binding in metazoans, uncovering factors that facilitate ORC recruitment still remains an under-studied aspect of genome replication (Eaton et al., 2011; Shen et al., 2010). To identify factors that interact with ORC with the potential to facilitate ORC binding to chromatin or regulate ORC activity, we immunoprecipitated endogenously tagged ORC2-GFP from Drosophila embryo extracts (Fig. 1A). Importantly, extracts were treated with benzonase to ensure ORC2-associated proteins were not indirectly bridged by DNA. We used a stringent statistical cut off to define ORC2-associated proteins (p-value of less than 0.05 and a fold enrichment greater than 2, see Materials and Methods; Supplemental Table 1). Using these parameters, we identified all six subunits of the ORC complex (Fig. 1C). Surprisingly, we also identified six Nups (Elys, Nup98-96, Nup75, Nup160, Nup133, and Nup107) that were statistically enriched in ORC2-GFP immunoprecipitation (Fig. 1B; Supplemental Table 1). Interestingly, five out of the six ORC2-GFP-associated Nups are members of the Nup107-160 complex that form rings on the inner and outer faces of the nuclear pore (Beck and Hurt, 2017). Given that an antibody specific to Drosophila Elys was available, we used IP followed by Western blotting to independently validate the association between ORC2 and Elys (Fig. 1D).

**Figure 1.**
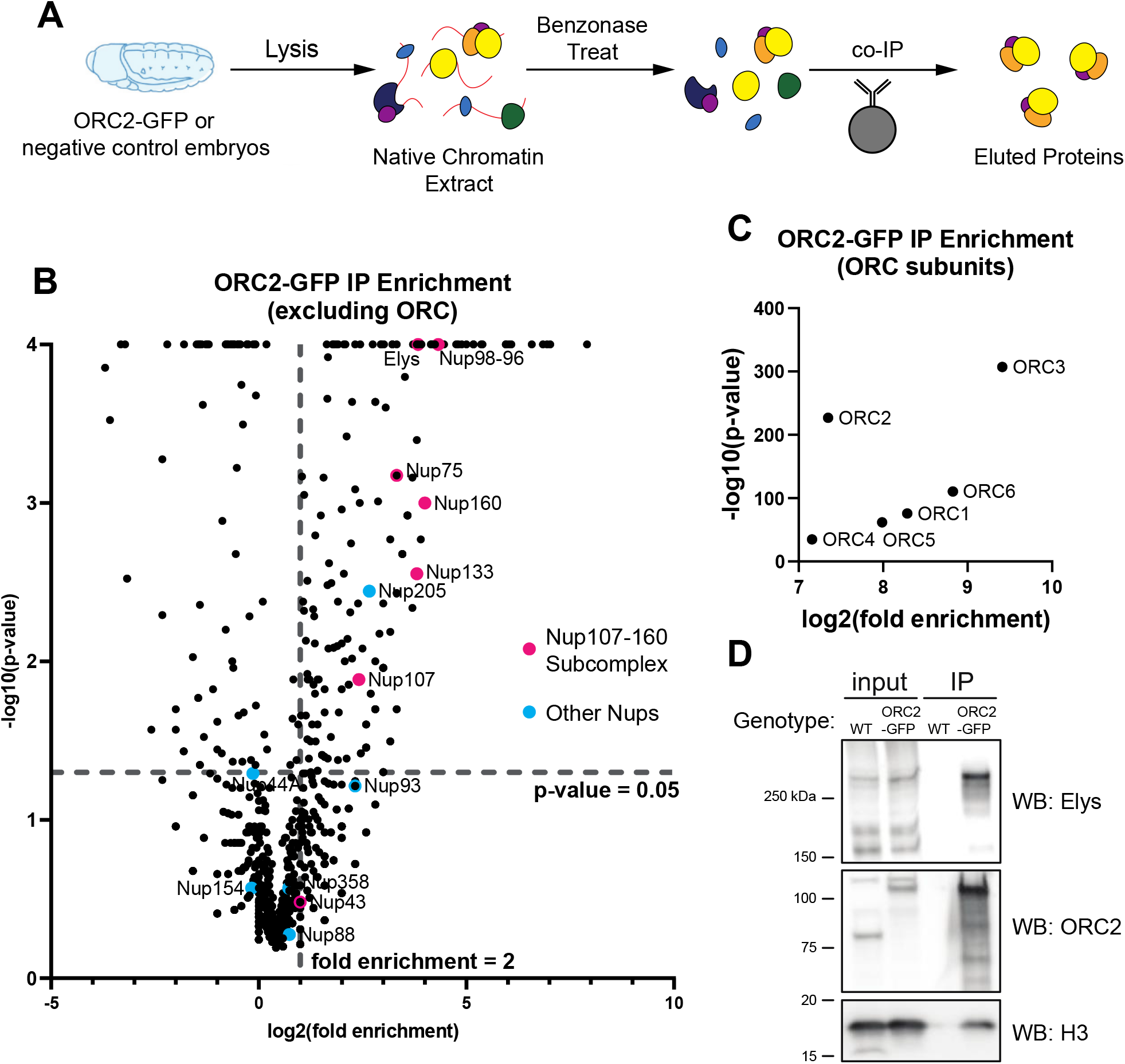
ORC interacts with subunits of the nuclear pore complex. (A) Schematic of extract preparation and immunoprecipitation protocol using *ORC2-GFP* or *Oregon R* (negative control) embryos. (B) Average fold enrichment and statistical significance for three biological replicates of GFP-Trap IP-mass spectrometry for *ORC2-GFP* expressing embryos relative to *OregonR* negative controls embryos. Log2 fold enrichment plotted against -log10 of p-value. Fold enrichment was calculated by dividing spectrum counts for GFP IP by the negative control. P-values were calculated by performing a Fisher’s Test for each individual protein. P-values less than 0.00010 were rounded for simplicity. Highlighted are all nucleoporin proteins identified by mass spectrometry. Dashed lines indicate significant level cut-offs (<0.05 for p-value and ≥2-fold enrichment). (C) Same as (B) but with only ORC subunits. (D) Western blots using anti-ORC2, anti-Elys, or anti-Histone H3 antibody on samples derived from the IP.

The two most enriched Nups identified, Elys and Nup98-96, have roles beyond being structural subunits of the nuclear pore (Kuhn et al., 2019; Panda et al., 2014; Pascual-Garcia et al., 2014, 2017). In Xenopus extracts, Elys associates with the activated replicative helicase, but not ORC (Gillespie et al., 2007). Furthermore, DNA replication is severely inhibited when Elys is depleted from extracts (Gillespie et al., 2007). Given that the Xenopus extract system more closely resembles early Drosophila embryogenesis, we repeated the ORC2 IPs throughout Drosophila embryogenesis to determine if the association between ORC2 and Elys was developmentally regulated. The association between ORC and Elys; however, occurred at multiple time points through embryogenesis and mirrored protein levels (Supp. Fig. 1B). Taken together we conclude that ORC2 associates Elys and several nucleoporins that make up the Nup107-160 subcomplex of the NPC.

### ORC2 binds the same genomic regions as several Nups

Given that ORC associates with Nups, we asked if ORC and Nups co-localize on chromatin. Nups bind to distinct regions of chromatin to regulate transcription and promote chromatin accessibility (Kuhn and Capelson, 2019). This aspect of Nup function is likely independent of nuclear pore function as individual Nups have distinct chromatin binding profiles (Capelson et al., 2010; Kalverda et al., 2010; Kuhn et al., 2019; Panda et al., 2014; Pascual-Garcia et al., 2014, 2017; Raices and D’Angelo, 2017; Vaquerizas et al., 2010). Using previously published ChIP-seq data sets generated in Drosophila S2 cells, we visualized the genomic binding profiles of ORC2 (Eaton et al., 2011) and multiple Nups representing distinct subcomplexes of the nuclear pore (Gozalo et al., 2020; Pascual-Garcia et al., 2017). We also performed CUT&RUN using the mab414 antibody, which recognizes FG repeats found in several Nups to determine the genomic binding sites of nuclear pores more broadly (Davis and Blobel, 1986). Qualitatively, the binding profile of ORC2 shows extensive overlap with the binding profiles of Elys, Nup107, Nup98 and mab414 (Fig. 2A). Next, we quantified ChIP-seq signal of Nups relative to ORC2 peaks and found that Nup and mab414 ChIP-seq or CUT&RUN signal was enriched within ORC2 peaks with Elys followed by Nup98 showing the strongest signal across all Nups (Fig. 2B). Strikingly, 98% of ORC2 peaks overlap with Elys binding sites (Supp. Fig. 2A). These data show that ORC2 and Nups bind many of the same genomic regions.

**Figure 2.**
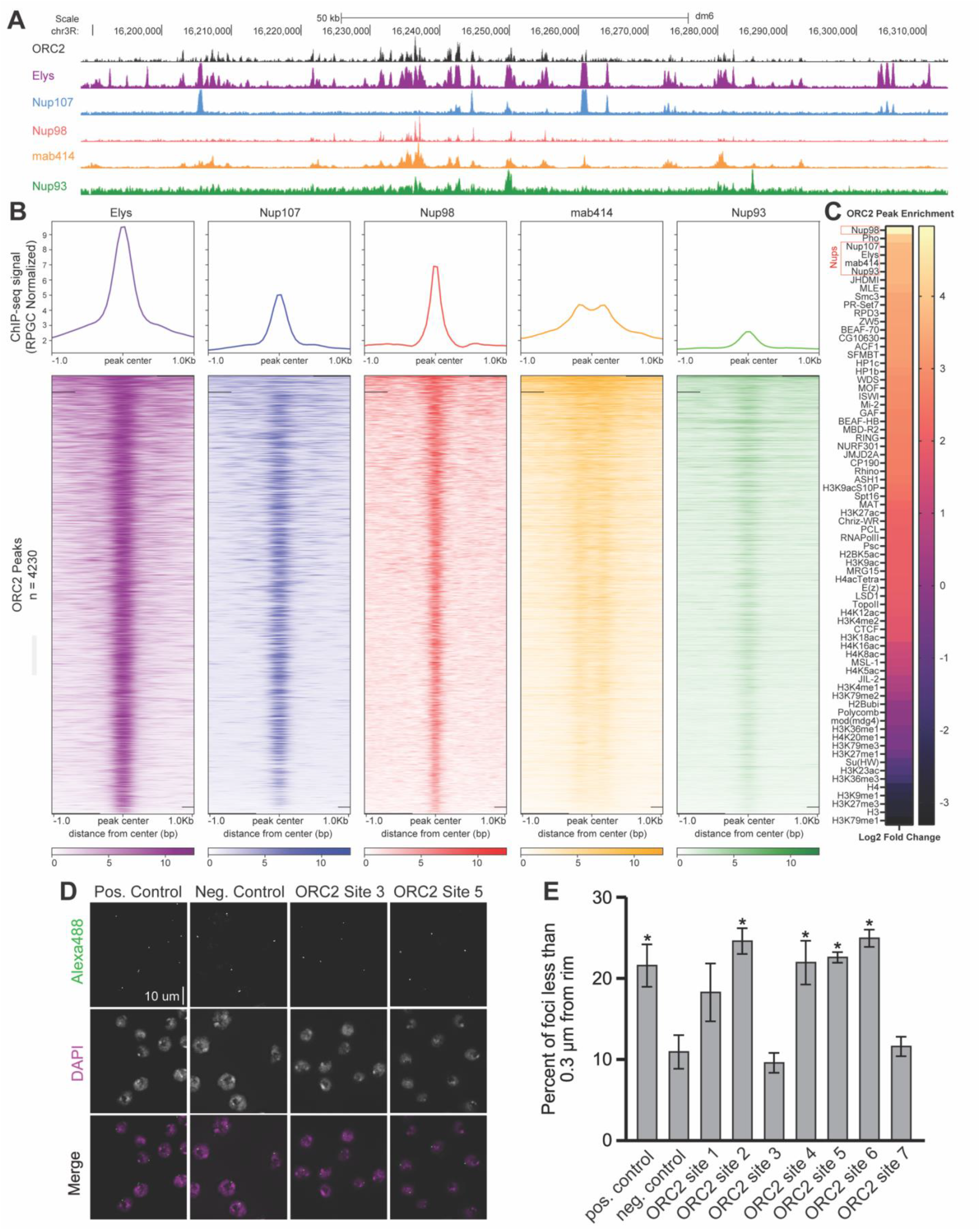
ORC2 binds the same genomic regions as several Nups. (A) Representative UCSC genome browser view of ORC2, Elys, Nup107, Nup98, mab414 and Nup93 ChIP-seq (or CUT&RUN) signal generated from previously published data. (B) Enrichment heatmap of ChIP-seq signals sorted by mean occupancy around the center of ORC2 peaks. (C) ORC2 peak enrichment heatmap for chromatin marks, transcription factors and Nup peaks from previously published data. Log2 fold enrichment for the observed overlap relative to the overlap expected from 1000 random permutations of the comparison peak set is shown. (D) Representative images of oligopaint performed in S2 cells for one positive (nuclear periphery associating) control site, one negative (non-nuclear periphery associating) control site and two ORC-binding sites that were also positive for Elys (Supp. Fig 2C for coordinates). (E) Quantification of the percent of oligopaint foci that were less than 0.3 μm from the nuclear rim for control sites and seven ORC-binding sites that are also positive for Elys. Asterisk indicates statistical significance relative to the negative control.

While we observed extensive overlap between ORC2 and Nup binding sites, we wanted to quantitatively measure the significance of this overlap relative to other chromatin-associated factors. To this end, we evaluated the overlap between ORC2 peaks, the available Nup ChIP-seq data sets, and all available S2 ChIP-seq data sets available from the modENCODE consortium. For each annotation, we compared the observed overlap with the overlap observed with 1000 randomly shuffled sets of peaks (Celniker et al., 2009; Contrino et al., 2012). This allowed us to test if the degree of overlap with ORC2 peaks was greater than the expected overlap if peaks were randomly distributed along the genome (see Materials and Methods). As a proof of principle, our analysis revealed several modENCODE factors that were either enriched or depleted at ORC2 binding sites consistent with previous work (Eaton et al., 2011). Strikingly, not only were Nups enriched at ORC2 binding sites, they were among the most statistically enriched factors (p-value = 0.0001, log2 fold change > 3.5 for Elys, mab414, Nup107, Nup93, Nup98) out of all 72 data sets we analyzed (Fig. 2C and Supp. Fig. 2B). Taken together, we conclude that Nup binding sites show significant overlap with ORC2 binding sites genome wide.

Nups are capable of binding chromatin independently of nuclear pores, but also bind chromatin when in complex with nuclear pores (Capelson et al., 2010; Kadota et al., 2020; Kuhn and Capelson, 2019). Given this, we were curious if the ORC2 binding sites that overlap with Nup binding sites required localization to the nuclear pore, suggesting the interaction between ORC and Nups occurs at NPCs. To formally test this, we selected seven 10-kilobase regions that were positive for both Elys and ORC2 binding (ORC2 sites 1-7) and generated oligopaint probes specific to these sites (Supp. Fig. 2B and 2C). We then measured the proximity of these seven sites to the nuclear periphery. If ORC binding and colocalization with Nups requires a functional nuclear pore, we would expect these sites to be enriched at the nuclear periphery. This, however, is not the case. Just over half of the sites we tested were found in close proximity to the nuclear rim (Fig. 2D and 2E). Given that ORC2/Elys binding sites are not required to be at the nuclear periphery, this suggests that the ORC/Nup association occurs independently of the nuclear pore.

### ORC binding to chromatin partially depends on Elys

So far, we have shown that ORC physically associates with the members of the Nup107-160 subcomplex and there is a high degree of colocalization between ORC and Nups on chromatin. To determine if there is a functional relationship between ORC and Nups, we asked if the chromatin association of ORC is dependent on Nups. To this end, we depleted either GFP (negative control), ORC2, Elys and Nup98-96 using RNA interference (RNAi) in Drosophila S2 cells. Nup98-96 was selected as a control as these genes are transcribed into a single mRNA, which is translated into a larger precursor protein that is ultimately cleaved to produce Nup98 and Nup96 (Fontoura et al., 1999). Therefore, RNAi against Nup98-96 reduces the steady state protein level of both Nup98 and Nup96 (Fontoura et al., 1999). Depletions were verified by Western blotting against Elys and ORC2 (Supp. Fig. 3A). We used Elys protein levels as a proxy for the Nup98-96 depletion since these proteins are in the same complex (Franz et al., 2007; Gillespie et al., 2007; Rasala et al., 2006; Shevelyov, 2020) and we consistently observed a reduction in Elys protein in the Nup98-96 depletion (Supp. Fig. 3A).

Given that Elys binds to chromatin to promote the decondensation of chromatin (Kuhn et al., 2019), we hypothesized that Elys, and perhaps other Nups, could promote ORC binding to chromatin, as ORC preferentially associates with open and active regions of chromatin (Eaton et al., 2011; MacAlpine et al., 2010). To test this, we quantified the amount of chromatin bound ORC2 in G1 phase nuclei in GFP, ORC2, Elys or Nup98-96 depletions using quantitative flow cytometry (Matson et al., 2017) (see Supp. Fig 3C for gating example) (Fig. 3A, 3B). G1 phase nuclei were selected because ORC is loaded in late M and G1, therefore, any differences in ORC chromatin association should be most apparent at this cell cycle stage. In ORC2-depleted control nuclei, we observed significantly less ORC2 on chromatin as expected. Consistent with our hypothesis, we observed significantly less chromatin-associated ORC2 in the Elys depletion relative to control cells (Fig. 3A, 3B). Interestingly, we do not observe a significantly different amount of chromatin bound ORC2 in Nup98-96 depleted nuclei, suggesting that not all Nups contribute to ORC loading onto chromatin (Fig. 3A, 3B). In fact, depleting Nup107 and Nup160 (both members of the Nup107-160 subcomplex) did not affect ORC2 chromatin loading, indicating that the reduction of chromatin bound ORC2 is not a generic defect of depleting Nups (Fig 3A, 3B, Suppl. Fig. 3E). As an important control, we performed the same experiment with a second set of dsRNAs against ORC2, Elys, and Nup98-96 to eliminate the possibility that our observations are due to nonspecific effects from the dsRNA (Supp. Fig. 3D). Taken together, we conclude that proper ORC2 chromatin association is dependent on Elys and that the reduction in ORC2 chromatin association is not a general defect caused by Nup depletion.

**Figure 3.**
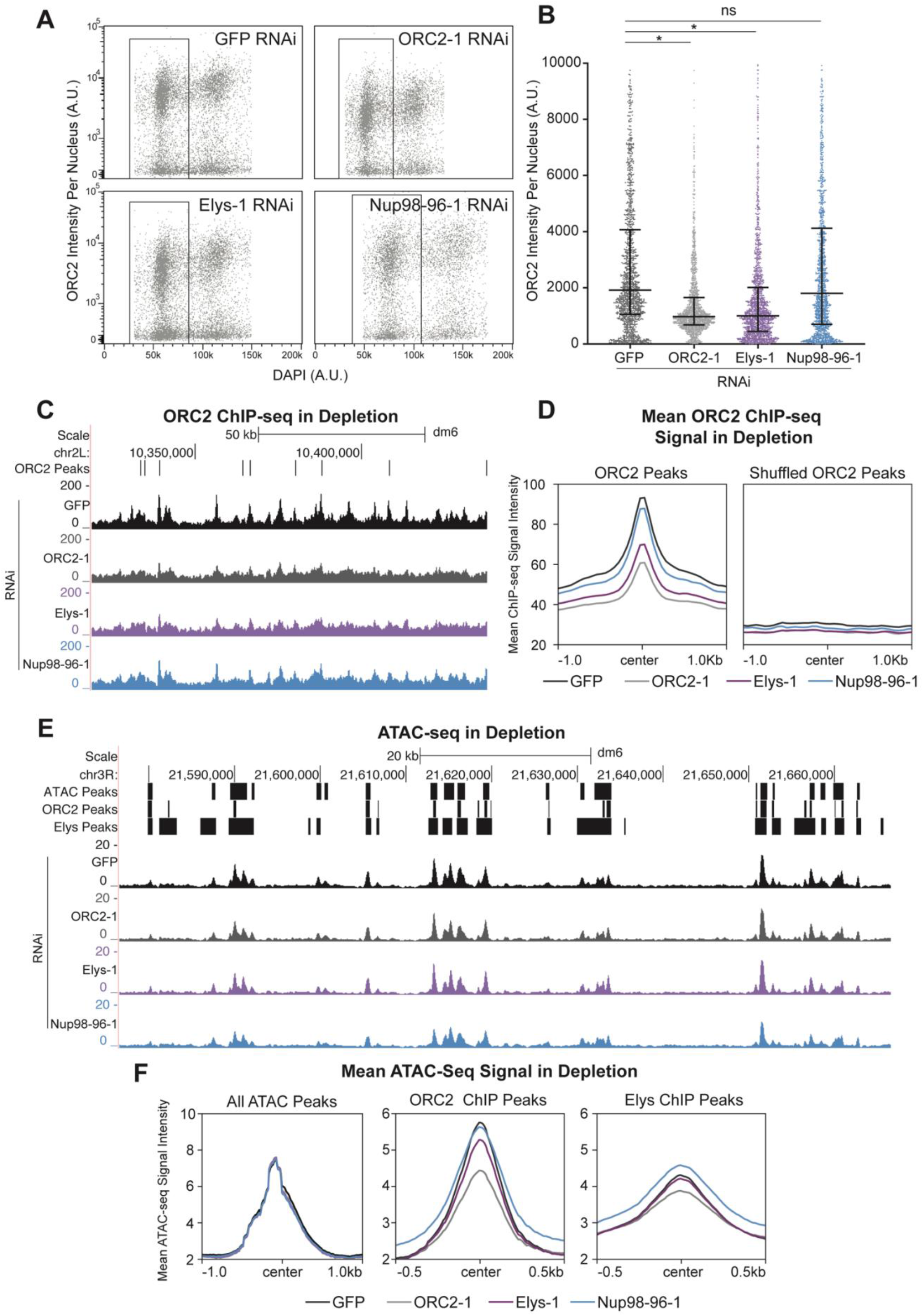
ORC’s chromatin association depends on Elys. (A) Horseshoe plot of nuclei with DNA content (DAPI) plotted against ORC2 intensity for each depletion from a single biological replicate. Black box indicates G1 population of nuclei used for the quantification in (B). A.U.: arbitrary units. (B) 1500 randomly selected G1 nuclei across three biological replicates were used to quantify the ORC2 intensity per nucleus for each depletion. Asterisk indicates p<0.0001 relative to the negative control by One-Way ANOVA with a post-hoc Dunnett’s test. NS: No Significance. (C) Representative UCSC genome browser view of ORC2 ChIP-seq profiles for each depletion. ORC2 binding sites (ORC2 Peaks - defined by Eaton et al., 2011) are indicated by black bars. (D) Quantification of mean ORC2 ChIP-seq signal for within defined ORC2 peaks or shuffled ORC2 peaks, centered on ORC2 peaks or shuffled ORC2 peaks, respectively. (E) Representative UCSC genome browser view of ATAC-seq for each depletion. ATAC-seq peaks, ORC2 ChIP-seq peaks and Elys ChIP-seq peaks are indicated by black bars. (F) Quantification of mean ATAC-seq signal for either all ATAC-seq peaks (n=12771), ORC2 ChIP-seq peaks (n=4280) or Elys ChIP-seq peaks (n=12048) centered on their respective peaks. Note the scales are different for all ATAC-seq peaks plots.

Next, we asked if the reduction of chromatin-bound ORC2 upon Elys depletion occurs throughout the genome or if only specific genomic regions or ORC2 binding sites are affected. To answer this, we performed ChIP-seq using an ORC2 antibody in Drosophila S2 cells that were depleted for either GFP, ORC2, Elys, or Nup98-96. We then quantified the ChIP-seq signal intensity within previously identified ORC2 binding sites throughout the genome (Eaton et al., 2011). For our positive control, we observed less ORC2 ChIP-seq signal in the ORC2 depletion relative to the GFP negative control (Fig. 3C & 3D). Consistent with our flow cytometry results, there was less ORC2 ChIP-seq signal in the Elys depletion, but not Nup98-96 depletion (Fig. 3C & 3D). Furthermore, we observed a reduction in ORC2 signal throughout the genome, indicating that depletion of Elys impacts all ORC2 binding sites. To ensure the reduction in ORC2 ChIP-seq signal was specifically within ORC2 peaks, and not a general trend throughout the genome, we shuffled all ORC2 peaks across the genome and found no difference in the mean ORC2 ChIP-seq signal (Fig. 3D). Therefore, the reduction of signal is specific to ORC2 binding sites (Fig. 3D). Taken together, we conclude that depleting Elys results in less ORC2 binding throughout the genome and that Elys, but not the other Nups tested, facilitates ORC2 loading onto chromatin.

ORC is known to bind to open and accessible regions of the genome (Eaton et al., 2011; MacAlpine et al., 2010). Given that Elys is known to promote chromatin decondensation, one possibility is that Elys facilitates ORC loading indirectly by promoting chromatin accessibility. To test this possibility, we performed ATAC-seq in RNAi-treated cells to measure chromatin accessibility within each depletion (Fig. 3E). Importantly, there was no global change in accessibility when comparing all ATAC-seq peaks throughout the genome upon Elys, ORC or Nup98-96 depletions (Fig. 3F). When comparing accessibly specifically within ORC2 binding sites, we noticed a modest reduction in ATAC-seq signal upon Elys depletion and an even more significant change in accessibility upon ORC2 depletion (Fig. 3F). ORC can directly promote chromatin accessibility at ORC binding sites (Eaton et al., 2010). Given the reduction in ORC2 binding upon Elys depletion, it is likely that the modest change in accessibility upon Elys depletion is an indirect effect of reduced ORC binding at those sites. Consistent with this, there was no significant difference in accessibility at Elys binding sites that do not overlap with ORC2 binding sites (Fig. 3F). Together, these data argue that the reduction in ORC chromatin association in an Elys depletion may not be driven by changes in chromatin accessibility.

### Nup depletion sensitizes cells to fork stalling

Given that ORC associates with Nups, co-localizes with Nups on chromatin, and that the association of ORC with chromatin is at least partially dependent on Elys, we wanted to ask if depletion of Elys and Nup98-96 affects cell cycle progression and/or genome stability. We reasoned that if ORC loading on chromatin was compromised, then we may observe a defect in S phase entry. Therefore, we pulsed cells with EdU and measured the fraction of cells in G1, S and G2/M based on their DNA content and EdU status by flow cytometry. In our ORC2 depletion that serves as a positive control, we saw a modest increase in G1 cells and reduction in S phase cells relative to the GFP negative control, consistent with a defect in S phase entry (Supp. Fig. 4A and 4B). The modest effect is expected since excess ORC is loaded onto chromatin to ensure sufficient replication start sites to complete DNA replication (Kawabata et al., 2011). Depletion of Elys, however, did not significantly alter the cell cycle profile relative to the negative control (Supp. Fig. 4A and 4B). Given the modest effect observed with the ORC2 depletion, and the level of ORC still associated with chromatin upon Elys depletion (Fig. 3A-D), this was not entirely unexpected. Depletion of Nup98-96, however, drastically reduced the fraction of cells in S phase while increasing the fraction of cells in G1 and G2/M (Supp. Fig. 4A and 4B). Depletion of Nup98-96 did not significantly affect the level of chromatin-bound ORC (Fig. 3A-D). Therefore, we conclude that Nup98-96 influences cell cycle progression independently of ORC chromatin association.

Excess replication start sites are not always essential during an unperturbed S phase, but become critical upon replication stress (Alver et al., 2014). This is largely due to the need to fire dormant replication origins to complete DNA synthesis when replication is perturbed (Doksani et al., 2009; Ge et al., 2007; Ibarra et al., 2008). Given that we have observed a reduction of chromatin bound ORC but no change in the percent of cells in S phase in an Elys depletion, we considered the possibility that a reduction in chromatin-bound ORC could lead to a defect in dormant origin firing. If there are insufficient dormant origins upon an Elys depletion, then Elys-depleted cells should be sensitive to replication fork inhibition as there are less origins available to rescue stalled replication forks (Alver et al., 2014; Doksani et al., 2009; Ge et al., 2007; Ibarra et al., 2008). Therefore, we treated cells depleted for GFP, ORC2, Elys or Nup98-96 with a low dose of aphidicolin and measured the level of γH2Av (the Drosophila equivalent of γH2Ax in mammals) by quantitative immunofluorescence. We chose a dose of aphidicolin that did not increase the level of DNA damage, as measured by γH2Av, in our negative control (GFP) (Fig. 4A and 4B). We found that depletion of Elys and Nup98-96 alone caused a modest increase in DNA damage (Fig. 4A and 4B). Upon aphidicolin treatment, however, there is a significant increase in the amount of DNA damage both relative to the negative control (Fig 4A, bottom panel and Fig. 4B, right) and relative to the untreated depletions (Fig. 4A, and Fig 4B, pink bars). We conclude that the sensitivity to aphidicolin we observe is consistent with the possibility that dormant origin firing is reduced in an Elys depletion.

**Figure 4.**
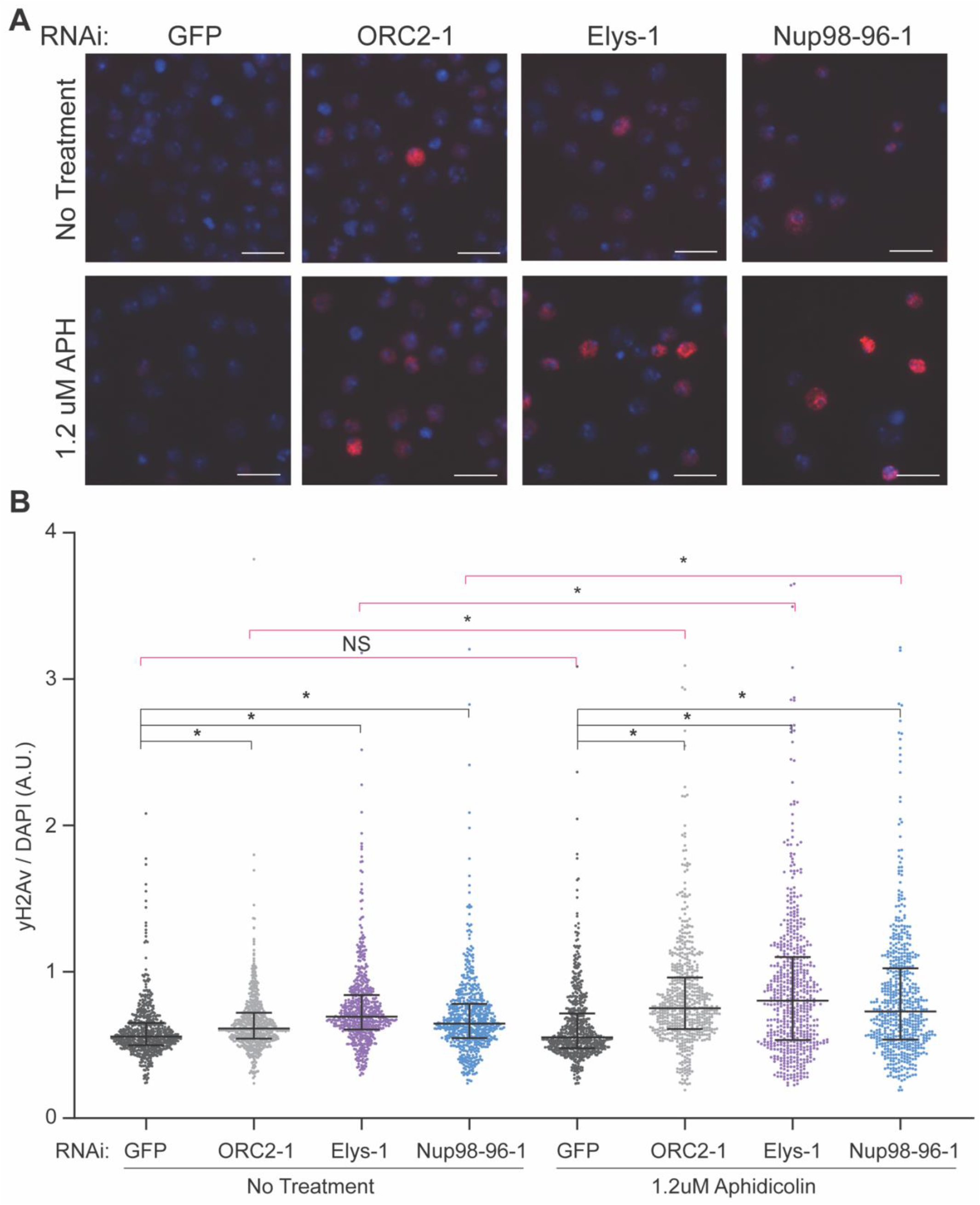
Nup depletion sensitizes cells to fork stalling. (A) Representative images of γH2Av immunofluorescence performed on RNAi-treated cells with or without aphidicolin treatment. Blue: DAPI. Red: γH2Av. Scale bar: 10 uM. (B) Quantification of (A). γH2Av and DAPI intensity for 600 total cells was quantified for each depletion with and without aphidicolin treatment. Each depletion contains 300 cells randomly selected from two biological replicates. Black bars indicate a One-Way ANOVA with a post-hoc Dunnett’s test comparing each depletion back to the negative control (GFP). Pink bars indicate a parametric T-test performed between each depletion comparing the untreated cells to the aphidicolin treated cells (GFP untreated vs. GFP treated for example). Asterisk denotes p < 0.0001. NS: No Significance.

## DISCUSSION

Our results show that ORC interacts with members of the Nup107-160 subcomplex of the nuclear pore, most notably the nucleoporins Elys and Nup98-96, establishing a link between replication inititation and Nups. Elys, Nup98, Nup93, Nup107 and FG-repeat-containing Nups are enriched at ORC2 binding sites and Nups are among the most significantly enriched chromatin factors at ORC2 binding sites. Strikingly, 98% of ORC sites are also Elys binding sites. It still unclear whether the interaction between ORC and Nups occurs on or off nuclear pores. Given that the ORC binding sites we visualized were not all localized to the nuclear periphery, this would suggest that the association between ORC and Nups are likely occuring off pore. Furthermore, if ORC and NPCs were present in the same protein complex, we would have expected to identify Nup subunits more broadly, rather than just a subset of Nups. Therefore our observations are most consistent with a model where Elys, and possibly other members of the Nup107-160 subcomplex, associate with ORC indpendently of the nuclear pore. This would be consistent with previously-published data where Elys and other Nups perform chromatin-related functions beyond their canonical role in the NPC.

Based on our present findings, we argue that Elys has a unique function in loading ORC onto chromatin. Importantly depletion of other Nups, including members of the Nup107-160 subcomplex, do not reduce the amount of ORC on chromatin. This reveals two important points. First, the reduction in chromatin-associated ORC upon Elys depletion is not a generic effect of altered NPC function. Second, out of the Nups tested, the ability to promote ORC loading seems to be unique to Elys. We do not rule out the possibility, however, that other Nups could contribute to ORC loading either independently or together with Elys. Interestingly, Elys and ORC both bind to chromatin in late M phase. It is possible that Elys, or another Nup, directly or indirectly interacts with ORC late in mitosis to facilitate ORC binding on chromatin by providing a molecuar bridge between chromatin remodelling and ORC. While we did not observe a global change in chromatin accessibilty upon Elys depletion, it is possible that Elys, together with it’s known interactor PBAP, could generate a nucleosome free region that would be optimal for ORC binding. If this happens specifically in late M phase, then it would be difficult measure changes in chromatin accessibilty by ATAC-seq from an asychronous population of cells.

Failure to load enough ORC throughout the genome could result in insufficent helicase loading. Given that the number and distribution of loaded helicases is necessary to maintain genome stability, depletion of Elys could compromise genome integrity due to a defect in origin licensing. Consistent with this, depletion of Elys shows an increased sensitivity to replication fork stalling. One possible explaination is that upon Elys depletion, there is insufficient ORC to promote dormant origin licensing upon for stalling. Interestingly, we observe a similar sensitivity to fork stalling in the Nup98-96 depletion. We predict this occurs through a different mechanism than the Elys depletion; however, as Nup98-86 depletion results in a stark reduction in cells in S phase and increase in cells in G2/M but does not significantly change ORC levels. Dormant origin usage could still underlie the sensitivity to fork stalling in Nup98-96 depleted cells. One possibility is that Nup98-96 affects helicase activation. This would explain why Nup98-96-depleted cells have a decrease in S phase and increase in G1 phase populations. Additionally, a failure to fire dormant replication origins would explain the increased sensitivity to fork stalling. Understanding how Nups differentially affect genome duplication and stability is an exciting area of future research.

## ACKNOWLEDGEMENTS

We thank Martina Ramos for expressing and purifying ORC2 and Elys for antibody generation. We thank the Vanderbilt Antibody and Protein Resource core for affinity purification of antibodies. The Vanderbilt Antibody and Protein Resource core is supported by the Vanderbilt Institute of Chemical Biology and the Vanderbilt Ingram Cancer Center (P30CA68485). Oligopainting was also supported from Vanderbilt-Ingram Cancer Center Support Grant (P30CA068485). Illumina sequencing was performed at the Vanderbilt VANTAGE core. We thank Hayes McDonald and the Vanderbilt Mass Spectrometry Resource Core for mass spectrometry. We thank Shelby Blythe and Maya Capelson or providing the Orc2-GFP flies and anti-Elys antibody used in an early phase of this work, respectively. We thank David Cortez, Katherine Friedman and Bill Tansey for providing feedback on the manuscript. This research was supported by the National Science Foundation grant (MCB-818019) and National Institutes of Health grant (R35GM128650) to J.T.N. and National Institutes of Health grant (1R35GM127087) to J.A.C. L.R. is supported by National Institutes of Health grant (1F31GM142286).

## AUTHOR CONTRIBUTIONS

**L.R**. Conceptualization, Formal Analysis, Investigation, Writing – Original Draft, Visualization; **C.L.L**. Formal Analysis, Investigation, Writing – Review & Editing; **M.L.B**. Formal Analysis, Writing – Review & Editing; **J.A.C**. Writing – Review and Editing, Funding acquisition; **J.T.N**. Conceptualization, Writing – Original Draft, Supervision, Funding acquisition

## DECLARATION OF INTERESTS

The authors have no competing interests

## MATERIALS AND METHODS

### Immunoprecipitations

Immunoprecipitations were performed on three biological replicates of both *ORC2-GFP* and *ORR* embryos. For each replicate, 0.5g of embryos aged 18-24 hours were collected, dechorionated, and flash frozen. Frozen embryos were ground thoroughly with a mortar and pestle in liquid N2. Ground embryos were then resuspended in NP40 Lysis Buffer (50 mM Tris HCl pH 8.0, 150 mM NaCl, 1% NP40, 1 mM EDTA, 1 mM EGTA, with 2X cOmplete™ Protease Inhibitor Cocktail EDTA-free (Millipore Sigma)). The embryonic extract was treated with benzonase nuclease (Millipore #7066) at a final concentration of 30 U/ml for 30 minutes at 4°C. After benzonase treatment, the extract was centrifuged at 4,000 rcf for 5 minutes. The supernatant was then used for the immunoprecipitation.

Prior to the immunoprecipitation, GFP Trap magnetic agarose beads (Chromotek #gtma-10) were washed and equilibrated with NP40 lysis buffer. Beads were added to extract and incubated for 2 hours at 4°C. After the 2 hours, beads were isolated and washed with 4 times with NP40 lysis buffer. Beads were then resuspended in 2x Laemmli sample buffer (Biorad #1610737) and boiled at 95°C for 20 minutes to elute protein.

### Mass Spectrometry

Eluates in Laemmli buffer were methanol/chloroform precipitated. After precipitation, immunoprecipitated samples were separated on a 4 – 12% NuPAGE Bis-Tris gel (Invitrogen), proteins were resolubolized in 5% SDS and prepared using S-trap (Protifi) using manufacturer’s protocol. Resulting peptides were desalted via C18 solid phase extraction and autosampled onto a 200 mm by 0.1 mm (Jupiter 3 micron, 300A), self-packed analytical column coupled directly to an Q-exactive+ mass spectrometer (ThermoFisher) using a nanoelectrospray source and resolved using an aqueous to organic gradient. Both the intact masses (MS) and fragmentation patters (MS/MS) of the peptides were collected in a data dependent manner utilizing dynamic exclusion to maximize depth of proteome coverage. Resulting peptide MS/MS spectral data were searched against the Drosophila protein database using SEQUEST (Yates et al., 1995). Identifications were filtered and collated at the protein level using Scaffold (Proteome Software).

Search results and peptide counts were refined in Scaffold using the following parameters: protein threshold false discovery rate = 5% minimum number of peptides ≥ 2, and a peptide threshold false discovery rate = 5%. Scaffold was used to perform a Fisher’s Test for each individual protein identified, comparing ORC2-GFP to the ORR negative control for all three replicates. For visualization purposes, p-values ≤ 0.0010 were rounded to 0.0010 in Fig. 2B. For Fig. 2C, p-values for ORC subunits were generated by a Fisher’s Test using R. Fold enrichment was calculated using the raw spectrum counts for individual proteins over the negative control. Volcano plots visualizing p-values and fold enrichment were made using GraphPad Prism.

### Western Blotting

The presence of ORC2 in the elute was confirmed prior to conducting mass spectrometry by SDS-PAGE followed by a Western blot for ORC2 using anti-ORC2 antibody. Briefly, samples were boiled and loaded onto a 4-15% Mini-PROTEAN TGX Stain-Free Gel (Biorad). After electrophoresis, the gel was activated and imaged using a BioRad ChemiDoc™ MP Imaging System following manufacturer recommendations. Protein was transferred to a low fluorescence PVDF membrane using a Trans-Blot Turbo Transfer System (BioRad). Membranes were blocked with 5% milk in TBST (140mM NaCl, 2.5mM KCl, 50 mM Tris HCl pH 7.4, 0.1% Tween-20). Blots were incubated with either anti-ORC2 antibody at 1:1000, anti-Elys antibody at 1:250, or HRP anti-Histone H3 antibody (abcam #ab21054) at 1:1000 overnight at 4°C. After primary antibody incubation, blots were washed, incubated secondary HRP antibody (Jackson Labs 711-035-150), washed once more and imaged.

### Antibody Generation

Full length ORC2 tagged with Maltose Binding Protein (MBP) was expressed and purified from *E. coli*. Briefly, ORC2 was cloned into the pLM302 expression vector. The expression construct was transformed into Rosetta 2 (DE3) cells (Novagen) and cultures were induced with IPTG and His-MBP-ORC2 was purified on Ni-NTA beads (BioRad). Purified protein was injected into rabbits for serum generation and collection (Cocalico Biologicals). For affinity purification, serum was first passed over an MBP column to deplete MBP-specific antibodies and the flow through fraction was passed over a column of MBP-ORC2 and eluted. An Elys-specific antibody was generated as previously described (Pascual-Garcia et al., 2017) using the same techniques described above.

### Oligopaint Fluorescent In Situ Hybridization (FISH)

Oligo pools were generated using the PaintSHOP application (Hershberg et al., 2021) from the Drosophila dm6 reference genome. A complete list of oligos can be found in Supplemental Table 1. Oligopaint probe production and FISH was performed largely as previously described (Nguyen and Joyce, 2019). Oligo pools were resuspended 50 μl of ddH_2_0 and 1 μl was used for an initial PCR amplification along with 2.5 μl of 10 μM forward (GCGTTAGGGTGCTTACGTC) and reverse (CACCTCCGTCTCTCACCT) primers, 25 μl 2X Q5 master mix, and 19 μl ddH_2_0 with 30 sec 98°C denaturation, 30 sec 55°C annealing, and 30 sec 72°C extension steps repeated 34 times. The PCR product was purified using a MagExtractor PCR & Gel clean up kit (Toyobo NPK-601) according to the manufacturer’s protocol and resuspended in 20 μl ddH_2_0. A secondary amplification was performed by mixing 1 μl of the first PCR product with 100 μl 2X Q5 master mix, 10 μl of 10 μM forward B and reverse B primers, and 79 μl ddH_2_0. The PCR was performed using the same program as described above, and subsequently purified as described and resuspended in 30 μl ddH_2_0. A Megascript T7 (ThermoFisher AM1334) reaction was performed by mixing 14 μl of the secondary PCR product with 4 μl of ATP, CTP, GTP, and UTP solutions and 4 μl reaction buffer, 2 μl RNase inhibitor, and 4 μl RT mix. The T7 reaction was incubated overnight at 37°C. A Maxima H Minus RT (ThermoFisher EP0752) reaction was setup by mixing 40 μl of the T7 reaction with 30 μl 100 uM forward B primer, 19.2 μl 100 mM dNTPs, 60 μl 5X RT buffer, 3 μl RNase inhibitor, 4 μl Maxima H Minus RT, and 143.8 μl ddH_2_0; this was incubated at 50°C for 3.5 hr. RNA was degraded by adding 150 μl 0.5M EDTA and 150 μl NaOH to the reaction, then heating at 95°C for 5 min. The DNA was cleaned and concentrated using a DNA Clean & Concentrator-100 kit (Zymo Research, D4029) and the DNA was resuspended in 90 μl ddH_2_0. The concentration ranged from 200-400 ng/μl. Pools were stored at -20°C until use.

For FISH experiments, S2 cells were concentrated and incubated in 100 μl Schnieder’s Drosophila media + 10% FBS on poly-lysine-coated slides for 1-2 hours in a humid chamber underneath a strip of parafilm the size of a cover slip. The media was then aspirated and the slides were incubated in freshly-prepared fixative solution (1X PBS and 4% paraformaldehyde) after transferring to coplin jars for 10 min at RT. Slides were washed in 1X PBS then incubated in freshly-prepared 0.5% Triton X-100 for 15 min at RT. Slides were rinsed with 1X PBS then dehydrated with successive incubations with 70%, 90%, and 100% ethanol for 2 min each at RT. Slides were then washed with 2X SSCT (2X SSC and 0.1% Tween-20) for 5 min. Next the slides were incubated with 2X SSCTF (2X SSC, 0.1% Tween-20, and 50% formamide) pre-heated to 90°C for 3 min in a new coplin jar that was also pre-heated. Slides were next incubated in 2X SSCTF at 60°C for 20 min (also pre-warmed). During this incubation, hybridization mix was prepared by vigorously mixing 300 μl formamide, 120 μl 50% dextran sulfate and 60 μl 20X SSC. 20 μl of this hybridization mix was then mixed with 4.5 μl oligo pool (at 200-300 ng/ μl) along with 0.5 μl 100 mM dNTPs, which was a sufficient quantity for one slide. Slides were dried for 5 min, then the hybridization mix + probe was added on top of the fixed cells. This was covered with a cover slip and sealed with rubber cement and dried for at least 20 min at RT. Slides were placed into a humid slide incubator and heated to 92°C for 3 min, then incubated overnight at 37°C. The next day cover slips were carefully removed and the slides were washed with 2X SSCT (pre-warmed) at 60°C for 15 min, then 2X SSCT for 10 min at RT, then 0.2X SSCT for 10 min at RT. New hybridization mix was prepared as before, and 120 μl was mixed with 29.75 μl ddH_2_0 and 0.25 μl 100 mM secondary fluorescent oligo probe (sequence of AAGCACCCTAACGCTTCACGATCCAT covalently linked to Alexa Fluor 488 dye), which was sufficient for 5 slides. 25 μl of this mix was then added on top of the fixed cells and sealed with a cover slip and rubber cement and the slides were incubated at RT for 1-2 hr. Cover slips were carefully removed and the slides were washed with 2X SSCT (pre-warmed) at 60°C for 15 min, then 2X SSCT for 10 min at RT, then 0.2X SSCT for 10 min at RT. 10-15 μl of Vectashield + DAPI (Vector Laboratories) was added on top of fixed cells, which were then sealed under a cover slip with nail polish.

### Cell culture

Drosophila S2 cells were provided by Drosophila Genomics Resource Center (DGRC). Cells were maintained following DGRC guidelines. Cells were grown in Schneider’s medium (Gibco 21720024) supplemented with 10% heat-inactivated fetal bovine serum (ThermoFisher A3840001) and penicillin-streptomycin (ThermoFisher 15140122). Cells were passaged every 3-5 days and maintained at a concentration of 3×10^6^-1×10^7^ cells/mL.

### RNA Interference

Cells were diluted to 1.5×10^6^ cells/mL in serum-free media. 20 μg of dsRNA generated using the T7 Transcription Kit (ThermoFisher AM1334) was incubated with cells for 45 minutes. A list of primers used to generate dsRNA can be found in Supplemental Table 2. After 45 minutes, serum-containing media was added to the RNAi-treated cells. Cells were then incubated for 5 days at 25°C. To confirm the depletion, 1 million cells were harvested and lysed in CSK buffer (10mM PIPES pH 7.0, 300 mM sucrose, 100 mM NaCl, 3 mM MgCl_2_ with with 2X cOmplete Protease Inhibitor Cocktail EDTA-free (Roche)) for 8 minutes. 2X Laemmli sample buffer (BioRad) was added to lysates and samples were incubated at 95°C for 5 mins. Depletions were confirmed by SDS-PAGE followed by western bloting for Elys, ORC2, and Histone H3 as previously described.

### Bioinformatics

Previously published data generated by ChIP-seq in Drosophila S2 cells was retrieved for ORC2 (Eaton et al., 2011). Elys, Nup107, Nup93, (Gozalo et al., 2020), Nup98 (Pascual-Garcia et al., 2017), mab414 data was generated by CUT&RUN (see CUT&RUN methods). Sequencing reads were aligned to dm6 with Bowtie2 (Langmead and Salzberg, 2012) using the pre-set --very sensitive-local. Duplicate reads were flagged after alignment with Picard: MarkDuplicates (Broad Institute) using Galaxy (Afgan et al., 2016). Coverage files were generated using Deeptools: BamCoverage (Ramírez et al., 2016) with the following options: 1X normalization, bin size = 50 bps, effective genome size = dm6. Genomic coverage was visualized using the UCSC Genome Browser (Kent et al., 2002) as shown in Fig. 2A. For peak comparisons, previously published peak files were used. For mab414, statistically significant peaks over an IgG negative control were called using MACS2 (Feng et al., 2012). Deeptools: plotHeatmap was used to generate the mean ChIP-seq signal plots and heatmaps centered on ORC2 peaks as shown in Fig. 2B.

The ATAC-Seq and ORC2 ChIP-seq data in Figure 3 was processed similar as above with minor differences. To generate the coverage plots for visualization, the ATAC-Seq data was normalized by CPM (counts per million) with a bin size = 50. To scale the ORC2 ChIP-seq data to the background signal, 25,000 genomic regions, each 250 base pairs long, were randomly selected. The total reads within the randomly selected regions for each depletion was determined and scaled down to the depletion with the fewest reads. The scaled coverage files were plotted in the UCSC Genome Browser for both Fig. 3C and 3E. For both ORC2 ChIP-seq and ATAC-seq, the mean signal was determined using Deeptools: plotProfile for each set of peaks. To generate shuffled ORC2 peaks, ORC2 peaks were randomly distributed across the genome, and the number of peaks and the length of each peak were kept the same using BedTools: ShuffleBed.

### CUT&RUN

CUT&RUN was performed using previously published methods (Ahmad and Spens, 2019; Skene and Henikoff, 2017; Skene et al., 2018). Briefly, 1 million S2 cells were harvested and spun down at 600 x g. Cells were washed with PBS and followed by wash buffer (20 mM HEPES pH 7.5, 150 mM NaCl, 0.1% BSA, with 2X cOmplete™ Protease Inhibitor Cocktail EDTA-free (Roche) and 0.6 mM Spermidine). Cells were attached to ConA beads in binding buffer (20 mM HEPES pH 7.9, 10 mM KCl, 1 mM CaCl2, 1 mM MnCl2) for 10 minutes. Cells were blocked and permeabilized in DBE buffer (wash buffer with 2mM EDTA and 0.05% digitonin) for 10 minutes. Cells were then incubated with 1μg of mab414 antibody (BioLegend) in DBE buffer at 4°C overnight.

After primary antibody incubation, cells were washed twice in DBE buffer. pA-MNase (gift from Kami Ahmad) was diluted 1:400 in DBE buffer and added to cells. pA-MNase was allowed to bind for one hour at room temperature. Cells were then washed twice with wash buffer and suspended in cleavage buffer (wash buffer with 2 mM CaCl_2_). DNA cleavage was carried out for 30 minutes on ice, then immediately stopped with stop buffer (170 mM NaCl, 20 mM EDTA, 4 mM EGTA). Supernatant containing the cleaved DNA was collected from the cells and treated with RNAse A and Proteinase K. SPRIselect beads (Beckman Coulter) were used to purify the fragmented DNA. To prepare this DNA for sequencing, the NEBNext Ultra II DNA Prep Kit for Illumina (New England Biolabs) was used using according to the manufacturer guidelines and then sequenced using an Illumina NovaSeq6000 for 150bp PE reads.

### Random Permutation Analysis

Peaks were downloaded for histone modification and transcription factor binding sites identified by ChIP-chip or ChIP-seq in Drosophila from modENCODE (Celniker et al., 2009; Contrino et al., 2012). All available ChIP-seq data in S2 cells were considered in addition to previously published ORC2 (Eaton et al., 2011) and nucleoporin peaks (Gozalo et al., 2020; Pascual-Garcia et al., 2017). For each ChIP-seq factor, the amount of base-pair overlap was calculated between the given factor and ORC2 peaks. A permutation-based technique was used to determine whether the observed amount of overlap was more or less than expected by chance. Briefly, an empirical p-value was calculated for the observed amount of overlap by comparing to a null distribution obtained by randomly shuffling regions throughout the genome and calculating the amount of overlap in each permutation. The p-values were adjusted for multiple testing using the Bonferroni correction. In this analysis, the location of the ORC2 peaks was maintained and the locations of the histone modification or transcription factor binding peaks were shuffled. The length distribution of the shuffled peaks was matched to the original set, and excluded all gap and ENCODE blacklisted regions from consideration. 1000 permutations were performed for each marker and ORC2 pair.

### ChIP-Seq

ORC2 ChIP-seq was performed as previously described (MacAlpine et al., 2010). Briefly, 20 million S2 cells for each depletion were harvested and centrifuged at 600 rcf for 5 mins. Cells were washed twice with PBS and fixed for 10 minutes with 1% PFA at room temperature. Crosslinking was quenched by adding glycine to a final concentration of 125 mM and incubating at room temperature for 5 minutes. Cells were spun down and resuspended in RIPA buffer (50 mM Tris-HCl, 140 mM NaCl, 1 mM EDTA, 1% NP-40, 0.1% Na-Deoxycholate, 0.1% SDS with 2X cOmplete™ Protease Inhibitor Cocktail EDTA-free). Cells were incubated for 1 hour at 4°C and sonicated using a Diagenode Bioruptor for 4 rounds of 10 cycles (each cycle was 30 seconds on, 30 seconds off at max power). After sonication, chromatin extract was cleared by centrifuging at 21,000 rcf for 5 mins. The remaining supernatant was used as input for the chromatin immunoprecipitation.

After preparing the chromatin extract, 1μg of anti-ORC2 antibody was added and allowed to incubate for 2 hours at 4°C. Protein A beads were washed with RIPA buffer, added to the extract and incubated for one hour at 4°C. Beads were then washed twice with RIPA buffer, twice with high-salt RIPA buffer (500mM NaCl), once more with RIPA buffer and once with TE Buffer. To elute protein, beads were incubated with elution buffer (50 mM Tris-HCl pH 8.0, 10 mM EDTA, 1% SDS) at 65°C for 15 minutes. Protein-DNA cross links were reversed by incubating at 65°C overnight. To recover DNA, samples were RNase A and Proteinase K treated, and phenol:chloroform extracted. Next, the DNA was isopropanol precipitated. Once the DNA was purified, the NEBNext Ultra II DNA Prep Kit for Illumina (New England Biolabs) was used to prepare the samples for next-generation sequencing. Barcoded libraries were sequenced using an Illumina NovaSeq for 150bp PE reads.

### ATAC-Seq

For each depletion, 50,000 cells were harvested and washed with PBS. ATAC-seq was performed as described previously (Buenrostro et al., 2015) using an ATAC-seq kit (Active Motif) as described by the manufacturer. Briefly, cells were resuspended in cold lysis buffer and centrifuged at 1000xg for 10 minutes at 4°C. The cell pellet was resuspended in tagmentation buffer and the reaction was incubated at 37°C for 30 minutes. Tagmented DNA was purified and used to generate sequencing libraries following manufacturer’s protocol. Libraries were sequenced with an Illumina NovaSeq6000.

### Flow Cytometry

To generate cell cycle profiles for RNAi-treated cells, 10 million cells were first pulsed with 20 mM EdU for 20 minutes after five days of RNAi treatment. Next, cells were washed twice with PBS and fixed overnight in ice-cold 70% ethanol. After fixation, cells were again washed with PBS and permeabilized for one hour at room temperature with PBX (PBS with 0.1% Triton X-100). Incorporated EdU was click-labeled with an Alexa Fluor 555 Azide (Invitrogen). Once clicked labeled, cells were washed twice with PBX and DAPI stained overnight. For the cell cycle analysis in Suppl. Fig 4B, three biological replicates were performed and the percent of cells in each phase of the cell cycle was quantified.

To quantify the amount of chromatin bound ORC in nuclei, 50 million cells were harvested after each depletion. The protocol was adapted from Matson et al., 2017. Cells were thoroughly washed with PBS and then lysed in cold CSK buffer supplemented with 0.5% Triton X-100 and 2X cOmplete™ EDTA-free Protease Inhibitor Cocktail for eight minutes on ice. PBS with 1% BSA was added to lysates and nuclei were pelleted by centrifugation at 2000xg for three minutes. Nuclei were then fixed with 4% PFA in PBS for 15 minutes at room temperature. After fixation, PBS with 1% BSA was added and fixed nuclei were pelleted by centrifugation at 2000xg for 7 minutes. Nuclei were washed once with PBS supplemented with 1% BSA and 0.1% NP40 (Blocking Buffer). Nuclei were incubated overnight at 4°C with anti-ORC2 antibody diluted 1:200 in blocking buffer. After the primary antibody incubation, nuclei were washed with blocking buffer and then incubated with anti-rabbit antibody conjugated to Alexa Fluorophore 568 (ThermoFisher) diluted 1:500 for two hours at room temperature. Nuclei were then washed twice with blocking buffer and DAPI stained overnight.

DNA content, EdU intensity and ORC2 intensity were determined using a BD LSRII flow cytometer. Flow cytometry data was analyzed and plotted using FlowJo (BD Biosciences). For an example of gating for these experiments, see Supplemental Figure 3C. For quantifying the ORC2 intensity per nuclei for, 500 nuclei from three replicates were randomly selected and pooled for a total of 1500 nuclei for each depletion. To determine statistical significance, a one-way ANOVA was performed with a post-hoc Dunnett’s test comparing each depletion to the negative control (GFP).

### Immunofluorescence

After four days of RNAi treatment, 1-3 million cells were treated for 24 hours with 1.2 uM aphidicolin in PBS (Millipore Sigma cat#: A0781). Cells were attached to Concanavalin A coated slides for one hour at room temperature. Cells were washed with PBS and then fixed with 4% PFA for 15 minutes and permeabilized with permeabilization solution (0.5% Triton X-100) for 8 minutes. After briefly rinsing in PBS, cells were blocked for 30 minutes in TBS with 0.1% Tween-20 (TBST) supplemented with 2% Normal Goat Serum (Sigma Aldrich). Histone H2AvD phosphoS137 antibody (Rockland cat #: 600-401-914) was diluted 1:50 in TBST and incubated overnight at 4°C. Next, cells were washed three times with TBST for 5 minutes each and incubated with Alexa fluorophore 568-conjugated anti-rabbit secondary (ThermoFisher cat#: A-11011), diluted 1:200 for one hour at room temperature. Cells were washed thrice in TBST, DAPI stained and mounted with Vectashield.

For each biological replicate, slides for each depletion were imaged at 40X with the same intensity and exposure time for each channel. To quantify the γH2Av signal, Nikon’s NIS Elements software was used to generate regions of interests (ROIs) using DAPI to define the ROI. Mean TxRed (γH2Av) and DAPI intensity for each ROI was determined for 300 cells per replicate (600 cells total). To account for differences in DNA content, γH2Av intensity was normalized to DAPI intensity. A one-way ANOVA with a post-hoc Dunnett’s test was performed for either the untreated group or the treated group (1.2μM aphidicolin). To determine the effect of treatment within each depletion, a parametric T-test was performed.

## Notes

### Competing Interest Statement

The authors have declared no competing interest.

