## Supplemental Information for "Nucleoporins facilitate ORC loading onto chromatin"

**A**

| | | ORC2-GFP | | | negative control | | | Comparison of abundance* (p-value) |
|---|---|---|---|---|---|---|---|---|
| | | Rep. #1 | Rep. #2 | Rep. #3 | Rep. #1 | Rep. #2 | Rep. #3 | |
| **ORC** | **ORC2** | 316 | 339 | 323 | 4 | 0 | 2 | < 0.00010 |
| | ORC1 | 109 | 80 | 123 | 2 | 0 | 0 | < 0.00010 |
| | ORC3 | 463 | 446 | 452 | 0 | 0 | 0 | < 0.00010 |
| | ORC4 | 58 | 43 | 42 | 0 | 0 | 0 | < 0.00010 |
| | ORC5 | 113 | 72 | 69 | 0 | 0 | 0 | < 0.00010 |
| | ORC6 | 148 | 151 | 155 | 0 | 0 | 0 | < 0.00010 |
| **Nup107-160 Subcomplex** | Elys | 16 | 31 | 96 | 6 | 2 | 2 | < 0.00010 |
| | Nup96-98 | 5 | 6 | 9 | 0 | 0 | 0 | < 0.00010 |
| | Nup75 | 4 | 6 | 10 | 1 | 0 | 1 | 0.00067 |
| | Nup160 | 4 | 5 | 7 | 0 | 1 | 0 | 0.0010 |
| | Nup133 | 3 | 4 | 7 | 0 | 0 | 1 | 0.0028 |
| | Nup107 | 6 | 4 | 6 | 1 | 1 | 1 | 0.013 |
| | Nup44A | 20 | 12 | 7 | 27 | 7 | 9 | 0.051 |
| | Sec13 | 7 | 4 | 5 | 8 | 5 | 4 | 0.20 |
| | Nup43 | 0 | 0 | 2 | 0 | 0 | 0 | 0.33 |
| | Nup37 | 0 | 0 | 0 | 0 | 0 | 0 | n.a. |
| **Othr Nucleoporins** | Nup205 | 9 | 3 | 7 | 0 | 0 | 0 | 0.0036 |
| | Nup93 | 1 | 1 | 3 | 0 | 0 | 0 | 0.061 |
| | Nup358 | 6 | 11 | 16 | 4 | 8 | 8 | 0.27 |
| | Nup154 | 3 | 3 | 2 | 3 | 0 | 6 | 0.27 |
| | Nup88 | 2 | 1 | 2 | 0 | 0 | 3 | 0.53 |

*Fisher's Exact Test

**B**

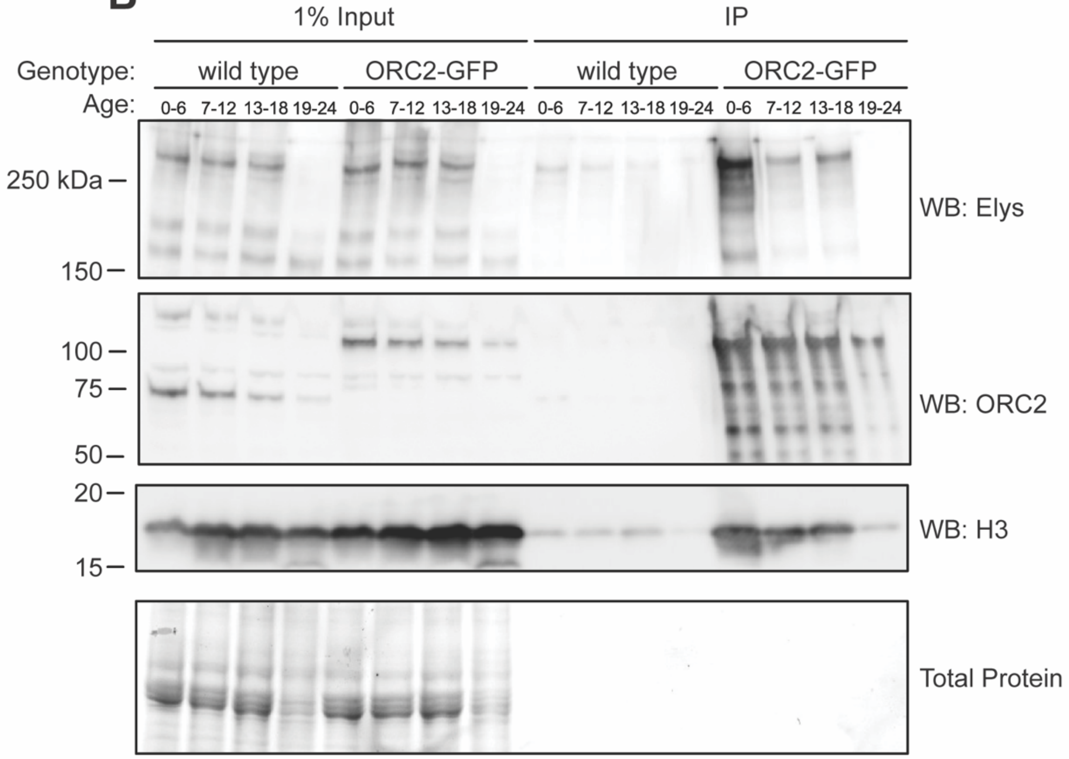

**Supplemental Figure 1. ORC2-GFP immunoprecipitation enriches for components of the Nup107-160 subcomplex of the nuclear pore.** (A) Table with peptide counts for three biological replicates of anti-GFP IP mass spectrometry done in either ORC2-GFP or negative control embryos as in Figure 1. Embryos were aged 16-24 hours. P-value was calculated by performing a Fisher’s Test. (B) Western blot of anti-GFP IP done in ORC2-GFP embryos throughout embryonic development. Western blots done using anti-Elys, anti-ORC2, or anti-Histone H3 antibodies. IPs were performed on embryos from indicated ages after egg laying (in hours). Total protein loaded for each sample shown in bottom box.

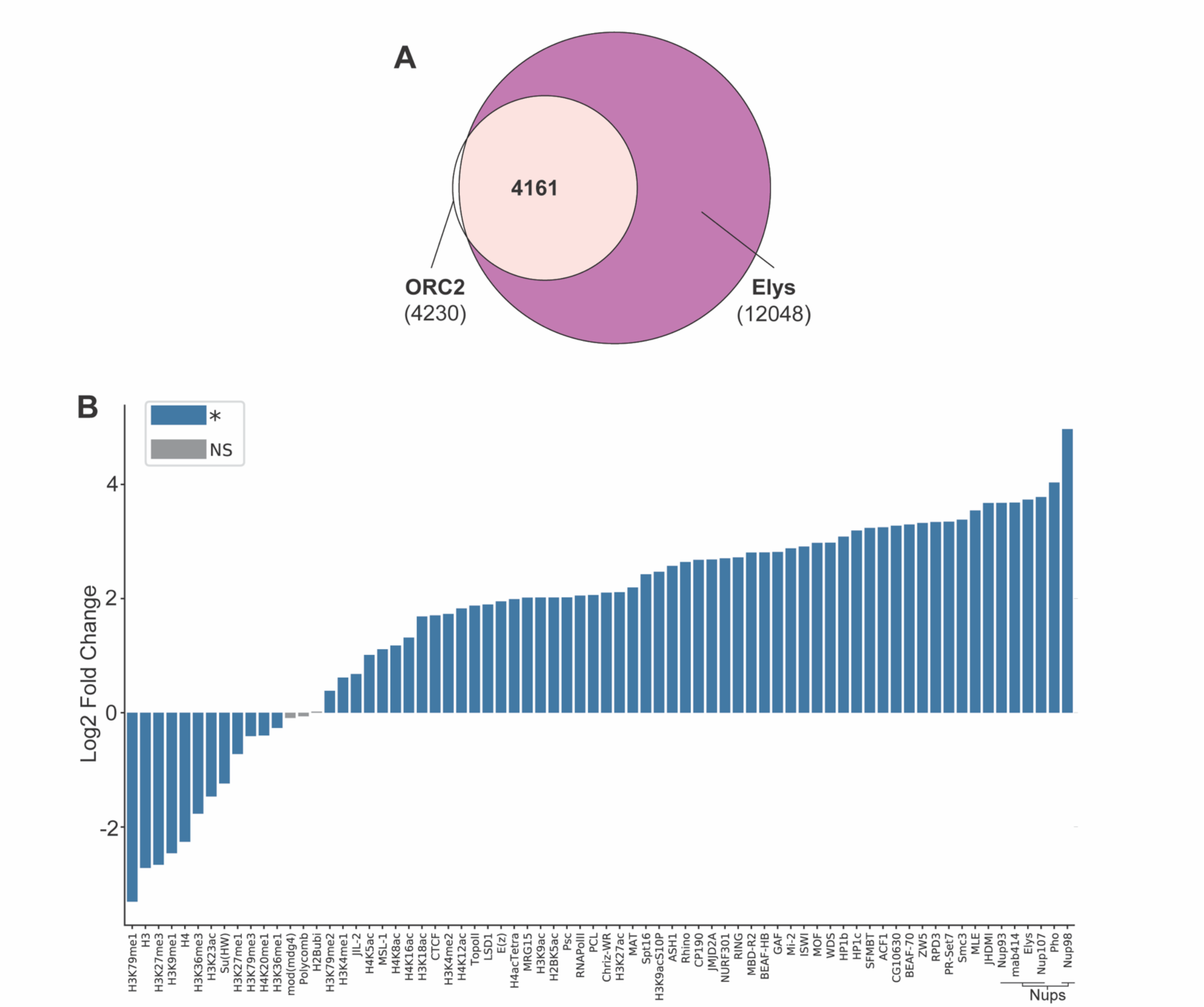


C

| Site Name | Coordinates |
|---|---|
| Pos. Control | chr2L:1352742-1360992 |
| Neg. Control | chr3R:19335000-19345000 |
| ORC2 Site 1 | chr3R:26879000-26888250 |
| ORC2 Site 2 | chr3L:14405500-14415500 |
| ORC2 Site 3 | chr3R:13644500-13654500 |
| ORC2 Site 4 | chr2L:12000000-12008000 |
| ORC2 Site 5 | chr2R:11600000-11611000 |
| ORC2 Site 6 | chrX:12727000-12742000 |
| ORC2 Site 7 | chrX:13822000-13830500 |

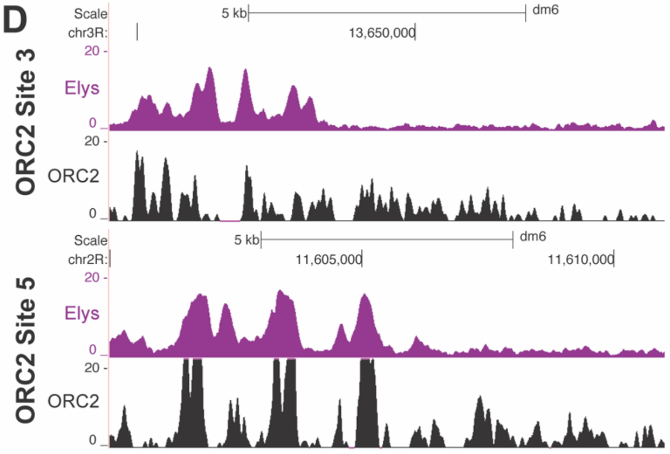


**Supplemental Figure 2. Nups are enriched at ORC2 binding sites.** (A) Venn diagram visualizing peak overlap between ORC2 (white) and Elys (purple). Number in parenthesis is the total number of peaks. Bold number is the number of ORC2 peaks that overlap with Elys peaks (4161 out of 4230). (B) Bar graph visualization depicting the log2 fold enrichments for data in Fig. 2C. Blue bars denote chromatin marks, transcription factors or nucleoporins that had a statistically significance correlation (positive or negative) with ORC2 peaks. Gray bars denote those that had a nonsignificant correlation. (C) Table containing site names and genomic locations of oligopaint probes for Fig. 2. (D) Representative genome browser view of Elys and ORC2 binding sties used for probes in oligopainting visualized in Fig. 2D (ORC2 sites 3 and 5).

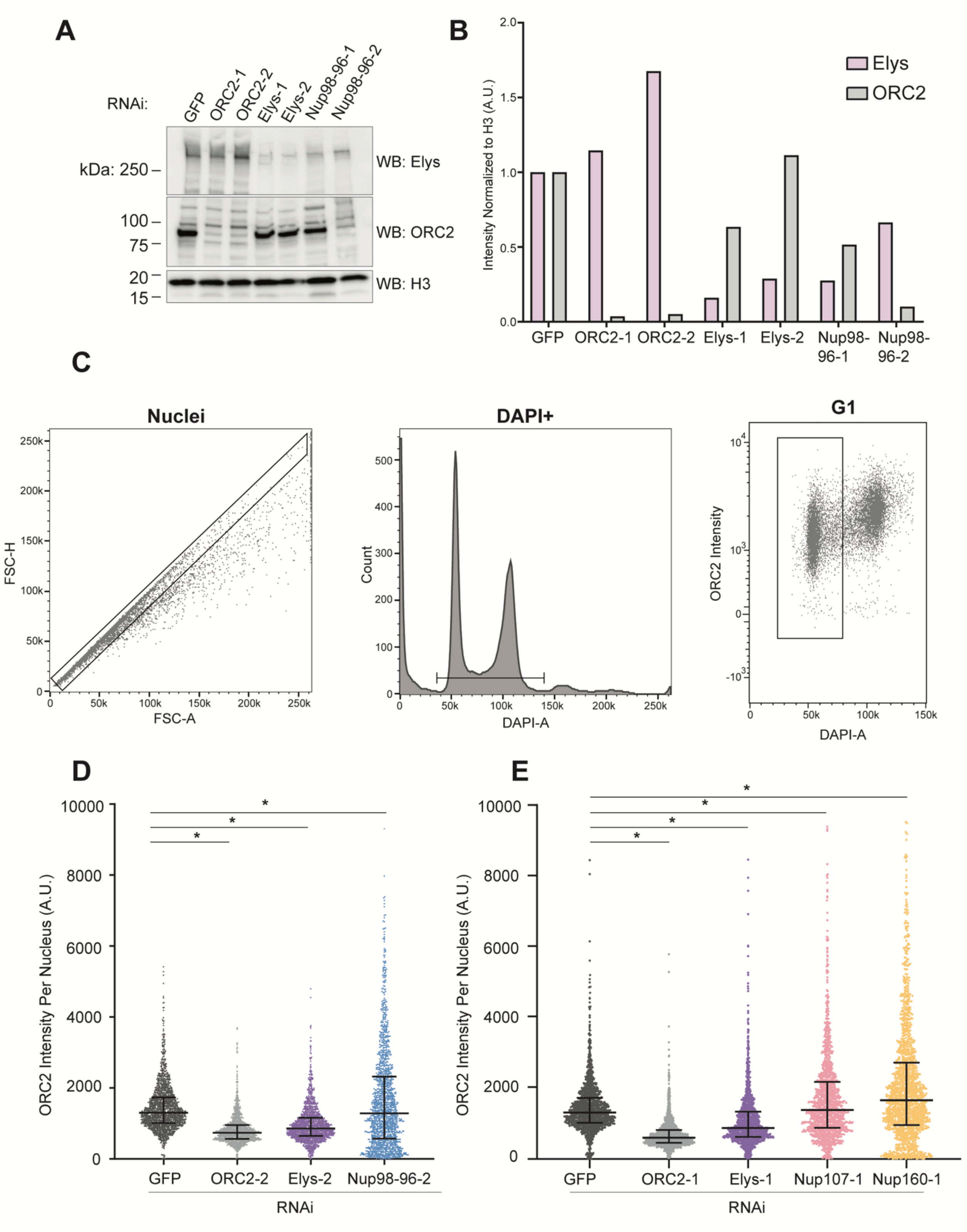


**Supplemental Figure 3. Not all nucleoporins contribute to ORC chromatin association.** (A) Western blot using anti-Elys, anti-ORC2 or anti-Histone H3 antibodies on samples prepared from cells treated with the indicated RNAi. (B) Quantification of (A). Western blot signal for Elys (pink) or ORC2 (gray) normalized to H3 for each depletion. (C) Gating example for G1 nuclei. Single nuclei isolated in the first gate indicated by the black box. DAPI positive nuclei were selected, indicated by black line, to generate a horseshoe plot. The first DAPI peak, shown as a black box in the third panel, was used for quantification. (D) Quantification of ORC2 intensity per nucleus using a second set of dsRNAs against ORC2, Elys, and Nup98-96. Each depletion shown contains 1500 nuclei taken from one biological replicate. Asterisk denotes a P value of $< 0.0001$ and was determined by a one-way ANOVA with a post-hoc Dunnett's test. (E) Same as (C) but with RNAi done using dsRNAs: GFP, ORC2-1, Elys-1, Nup107-1, and Nup160-1. Each depletion shown contains 750 nuclei randomly selected and pooled from two biological replicates for 1500 nuclei total.

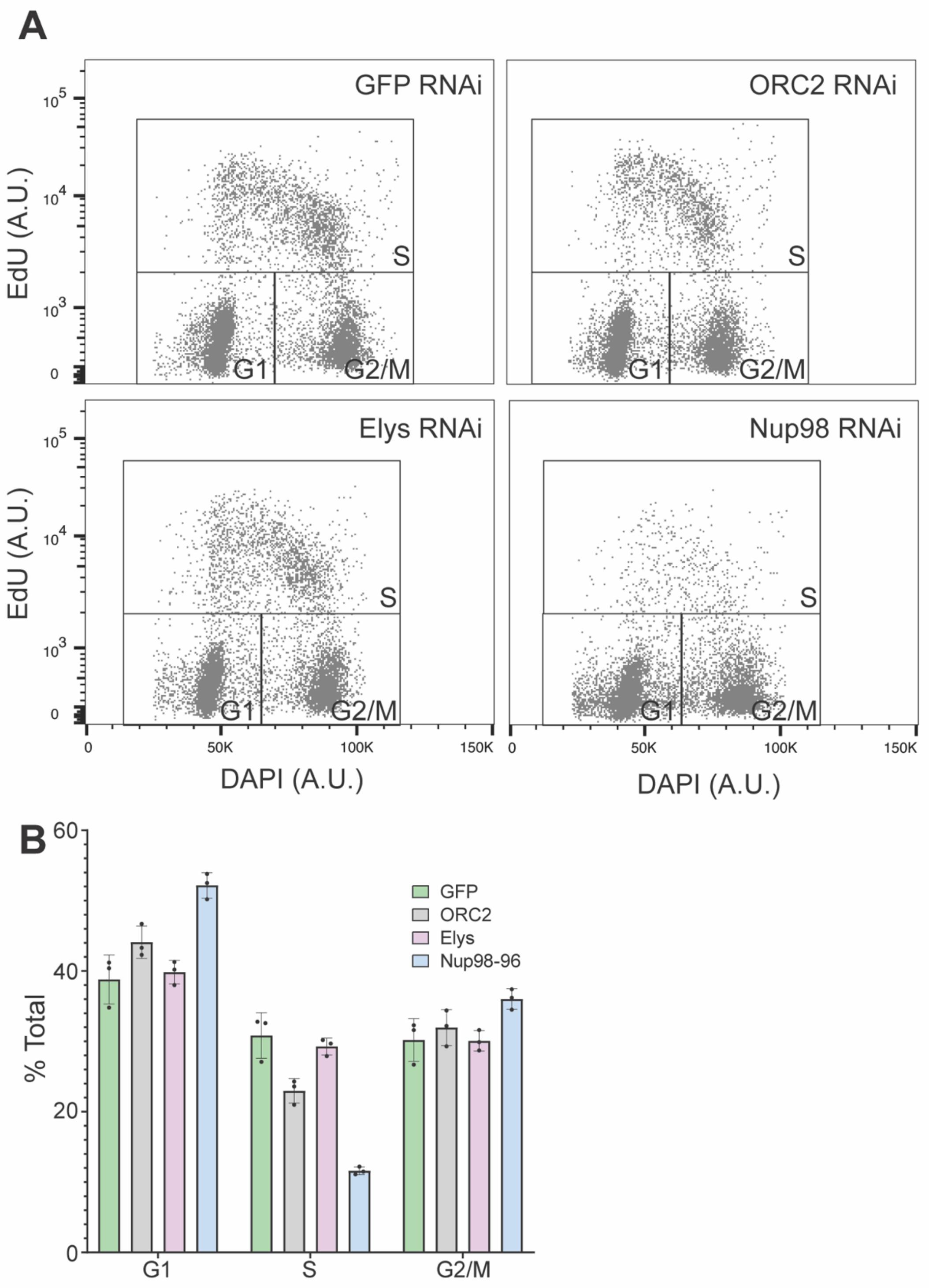


**Supplemental Figure 4. Nup depletions differentially affect cell cycle progression** (A) Horseshoe plot of RNAi-treated cells. Cells were DAPI stained and EdU pulsed to determine cell cycle phase. Black boxes indicate gating used to quantify percent of cell population within each cell cycle phase. (B) Percentage of cells in each indicated phase of the cell cycle (A). Shown are three biological replicates. Error bars represent the standard error of the mean.

| RNAi | Forward Primer | Reverse Primer |
|---|---|---|
| Elys-1 | TAATACGACTCACTATAGGGGCACGTATCTTCGCATCAGA | TAATACGACTCACTATAGGGGACAAGGACGCTTATTGGGA |
| Elys-2 | TAATACGACTCACTATAGGGTGGAGCCCTACCAAAAGAC | TAATACGACTCACTATAGGGCGCCTGGAGGAAATTTGG |
| Nup98-96-1 | TAATACGACTCACTATAGGGGGTGTGGCACCAAAAAGAGT | TAATACGACTCACTATAGGGCACCAATGTTTTTGGCAGTG |
| Nup98-96-2 | TAATACGACTCACTATAGGGGGAAGACCCAACTACCCGTT | TAATACGACTCACTATAGGGGCCCATTGGTCAAGGTCTAA |
| ORC2-1 | TAATACGACTCACTATAGGGAGCGATGCTGGCAACTC | TAATACGACTCACTATAGGGTATCCAGCATATCCTTGATGG |
| ORC2-2 | TAATACGACTCACTATAGGGTCGCTTGTGATGCTATCCAG | TAATACGACTCACTATAGGGAGCAAGATCCTCACTTCGGA |
| Nup107-1 | TAATACGACTCACTATAGGGATGCAGTATAGTAGGCTATTAG | TAATACGACTCACTATAGGGCGACGGCGGGTGTCTT |
| Nup160-1 | TAATACGACTCACTATAGGGGGCGGTTCATCTGGATCAA | TAATACGACTCACTATAGGGGCTGTGGGATCGCTTTTAC |
| GFP | TAATACGACTCACTATAGGGATGCCACCTACGGCAAG | TAATACGACTCACTATAGGGGTTCTGCTGGTAGTGGTC |

**Supplemental Table 2.** Primers used to generate dsRNAs for depletions.
